# Quantitative Properties of the Creation and Activation of a Cell-Intrinsic Engram

**DOI:** 10.1101/2020.03.17.995258

**Authors:** C.R. Gallistel, Fredrik Johansson, Dan-Anders Jirenhed, Anders Rasmussen, Matthew Ricci, Germund Hesslow

## Abstract

The conditional pause in the spontaneous firing of the cerebellar Purkinje, which determines the timing of the conditional eyeblink response, is mediated by a cell-intrinsic engram (Johansson, et al. 2014) that encodes the interstimulus interval. Our trial-by-trial analysis of the pause parameters reveals that it consists of a single unusually long interspike interval, whose onset and offset latencies are stochastically independent scalar functions of the interstimulus interval. The coefficients of variation are comparable to those observed in the timing of the overt conditional eyeblink. The onsets of the long interspike interval are step changes; there is no prior build-up of inhibition. A single spike volley in the parallel fiber input triggers the read-out of the engram into the long interspike interval; subsequent volleys have no effect on the pause. The high spontaneous firing rate on which the one-interval firing pause supervenes is markedly non-stationary (Fano factors >> 1).

## Introduction

The neurobiological memory mechanism carries forward in time, in computationally accessible form, facts gleaned from experience, such as the duration of experienced intervals (Balsam, Drew, & Gallistel, 2010; Buonomano, 2017; Mattell & Della Valle, 2018; Savastano & Miller, 1998). The experimental literature on the neurobiology of memory is vast, but little of it focuses on how and where specified quantitative facts are encoded in specified neurons.

The cerebellar Purkinje cell is the location of a memory for a quantitative experiential fact, namely, the duration of the inter-stimulus interval in the classically conditioned eye blink (Jirenhed, Bengtsson, & Hesslow, 2007; Jirenhed, Rasmussen, Johansson, & Hesslow, 2017; Johansson, Carlsson, Rasmussen, Yeo, & Hesslow, 2015; Johansson et al., 2014). The behaviorally observed conditional blink is driven by a conditional pause response in the spontaneous firing of specific blink-controlling Purkinje cells. These cells are inhibitory on the cerebellar nuclei, so a pause in Purkinje cell firing translates into an excitatory output signal from the cerebellum. The conditional firing pause is the only cellular level associative learning phenomenon whose quantitative properties align with the behaviorally established properties of associative learning (Gallistel & Matzel, 2013; Jirenhed & Hesslow, 2016).

In a Pavlovian delay conditioning protocol, a neutral stimulus (the conditional stimulus or CS for short) is repeatedly presented at a short, fixed latency prior to an unconditional stimulus (US for short), which is a stimulus that directly elicits a reflexive response in the naive subject. In an eye blink conditioning protocol, the US is a threat to the eye. The CS is any of a wide variety of stimuli that do not elicit a blink prior to conditioning. When the US follows the CS over a number of trials at intervals in the range from 0.1 to as much as 2.0s, a conditional blink to the CS develops. The number of conditioning trials prior to its appearance varies from a few to several hundred, depending on the subject and the parameters of the protocol (Gallistel, Balsam, & Fairhurst, 2004; Gallistel & Gibbon, 2000).

The conditional blink to the CS commences prior to the onset of the US, and it occurs on probe trials, when there is no US. The latency at which the CS evokes the blink varies in proportion to the interval that elapses between CS onset and US onset, so that the closure of the lid or membrane peaks near the moment when the threat to the eye is anticipated (Kehoe, Olsen, Ludvig, & Sutton, 2009; White, Kehoe, Choi, & Moore, 2000). Thus, this simple Pavlovian conditioning procedure inscribes in the brain a simple quantitative fact—the duration of the CS-US interval. The inscribed quantitative fact is read out into an appropriately timed behavior whenever the CS is again presented. The locus of the material change in the brain that encodes this fact is a prime target in the search for the engram, the neurobiological basis of memory.

The CS-conditional blink of the eye has been obtained in decerebrate preparations of cats, rabbits, ferrets and guinea pigs (Hesslow & Ivarsson, 1994; Kelly, Zuo, & Bloedel, 1990; Kotani, Kawahara, & Kirino, 2002; Mauk & Thompson, 1987; Norman, Buchwald, & Villablanca, 1977), proving that the forebrain is not essential; the brain stem alone is sufficient. Within the brain stem, the cerebellum is known to be the main locus of the memory trace (Freeman, 2015; Hesslow & Yeo, 2002; McCormick & Thompson, 1984; Thompson & Steinmetz, 2009). Disruption of cerebellar afferent signaling by a cerebral-vascular accident prevented eye blink conditioning in a human subject (Solomon, Stowe, & Pendlbeury, 1989).

The cerebellar Purkinje cell is among the largest neurons in the vertebrate brain (Apps & Garwicz, 2005; Knierem, 1997-Present; Llinas, Walton, & Lang, 2004). It is the sole output of the cerebellar cortex. Its immense, flat, densely arborized dendritic tree straddles a subset of the dense projections of parallel fibers. The parallel fibers arise from the tiny granule cells in the granular layer of the cerebellar cortex. They project upward to the top layer of the cerebellar cortex, where they bifurcate and run parallel to the folds of the cerebellar cortex. They pass through and make glutamatergic synapses on the staggered dendritic trees of numerous Purkinje cells. On the order of 200,000 parallel fibers synapse on the dendrites of each Purkinje cell. The granule cells from which they arise constitute substantially more than 50% of the neurons in a vertebrate brain (D’Angelo et al., 2013).

Climbing fibers provide the only other excitatory input to Purkinje cells (Apps & Garwicz, 2005; Knierem, 1997-Present; Llinas et al., 2004). They arise from cells in the inferior olivary nucleus. In stark contrast to the parallel fibers, only one climbing fiber innervates a Purkinje cell, but its terminal arbor wraps the cell with a dense engulfing bush of synapses. Short bursts of high-frequency presynaptic spikes in the climbing fiber produce the complex, multi-modal post-synaptic spike, which is thought to control Purkinje cell spiking rates and induce learning.

The main site of learning seems to be in the cerebellar cortex (Heiney, Kim, Augustine, & Medina, 2014; Hesslow, 1994; Hesslow, 1994b; Johansson et al., 2016; Ten Brinke et al., 2017; Yeo et al., 1984; Yeo, Hardiman, & Glickstein, 1985a, 1985c; Yeo, Hardiman, & Glickstein, 1986), which consists of roughly 1,000 microzones (Apps & Garwicz, 2005). Each part of the body maps by way of the climbing fiber system to several disparately located microzones. The Purkinje cells whose conditional pauses we here analyze are located in the C3 zone of the ferret cerebellum, the area that has been shown to mediate the classically conditioned eye blink (Heiney et al., 2014; G. Hesslow, 1994a; G. Hesslow, 1994; Jirenhed & Hesslow, 2016; Yeo, Hardiman, & Glickstein, 1985b, 1986; Yeo, M.J. Hardiman, & Glickstein, 1985). Like most Purkinje cells, these have a high spontaneous firing rate—on the order of 40-80 Hz.

The conditional pause in Purkinje cell firing develops in the decerebrate ferret even when the CS is direct electrical stimulation of the parallel fibers and the US is direct electrical stimulation of the climbing fiber (Johansson et al., 2014). Stimulation of off-beam parallel fibers—fibers that do not synapse on the Purkinje cell from which one is recording—produces a profound inhibition of the spontaneous firing, presumably by way of the stellate cells or the basket cells or both. However, a dose of a GABA blocker sufficient to block this inhibitory effect does not block the elicitation of the conditional pause (Johansson et al., 2014). This result implies that the pause is not mediated by input from the inhibitory interneurons synapsing on the Purkinje cell. This conclusion is further strengthened by the fact that the ability of the glutamatergic input from the parallel fibers to trigger the pause in firing is blocked by the infusion of an agent that selectively blocks the mGlu7 receptor in the synapse that a parallel fiber makes onto a Purkinje cell (Johansson, 2015; Johansson et al., 2015). Together these results make a strong case that the mechanism that times the CS-US interval, the mechanism that records into memory the result of that timing (the duration of the interval) and the mechanism that reads the remembered durations out into a pause of corresponding duration are intrinsic to the Purkinje cell itself (Johansson, 2019).

Previous analyses of the Purkinje cell pause responses have been based on averaging across trials, such as peristimulus time histograms. Although an indispensable technique, averaging can conceal and distort important features of the individual responses. For instance, sudden response onsets with variable latencies will look like gradual onsets. We have therefore re-analyzed a large body of data and here report the trial-by-trial statistics of the conditional pauses obtained from decerebrate ferrets under three experimental conditions. In the first conditions, the CS was pulsatile electrical stimulation of the mossy fibers at 50 Hz. (The mossy fibers are the input to the granule cells.) The US was two short bursts of pulsatile electrical stimulation to the inferior olive or to a climbing fiber. In the second, the CS was pulsatile electrical stimulation of the dorsum of the forepaw at 50 Hz and the US was two short bursts of pulsatile electrical stimulation to a climbing fiber. In the third, the CS was pulsatile electrical stimulation of the parallel fibers, which are the immediate presynaptic input to the Purkinje cell; the US was again two short bursts of pulsatile electrical stimulation to a climbing fiber. The CS-US interval used in training varied from 0.15 to 0.45s. The CS stimulation terminated just before US onset in some cases, while in others, it continued well beyond US offset.

## Determining Trial-by-Trial Pause Statistics

The conditional pause is commonly visualized by a raster plot of a sequence of probe (that is, CS alone) trials (Figure 1a). Its average duration may be estimated from the peri-CS histogram of spike counts (Figure 1b). In this work, we determine the distribution of pause statistics for individual trials—onset latencies, offset latencies, pause widths, pause depths and the abruptness of pause onsets. In the Discussion, we consider how these statistics constrain possible mechanisms for the generation of the pauses.

**Figure 1.**
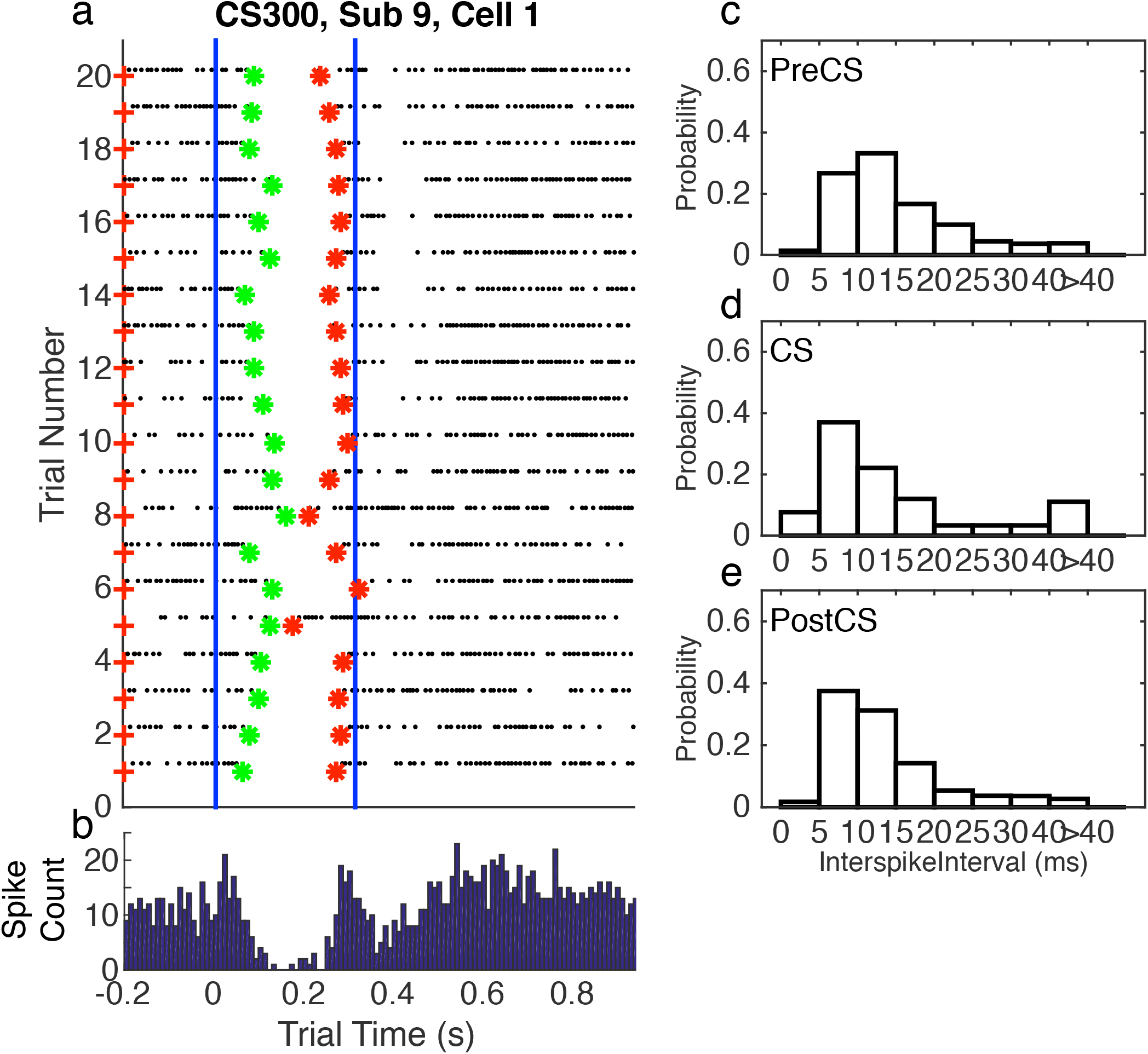
a. Spike raster plot of 20 successive probe trials (trials without a US) following conditioning with a 0.3 s CS-US interval. The CS was 0.3s stimulation of the dorsum of the forepaw at 50 Hz, both during training, when US onset immediately followed CS termination, and in these probe trials. The US during training was two 5-pulse bursts of 500 Hz stimulation of the climbing fiber input (5 ms between the bursts). The blue vertical lines mark the beginning and end of the CS-US interval. Green asterisks mark the pause onsets, as determined by our algorithm for estimating pause onsets and offsets; red asterisks mark the estimated pause offsets. b. Peri-CS histogram, counting spikes across the 20 trials into successive 10 ms wide bins. c-e. Probability distribution functions (normalized histograms, with 5 ms wide bins) on the inter-spike intervals during the 0.3s preceding the CS (c), during the CS-US interval (d), and during the 1 s after the CS (e)

The compilation of trial-by-trial pause statistics presupposes an algorithm for determining their onsets and offsets. We developed a Bayesian algorithm that uses the statistics from the peri-stimulus histogram to set prior probabilities on the locations of the onsets and offsets on individual trials and on the rates of firing before, during, and after the pause. It is described in the Appendix. The data and the Matlab™ code implementing our analyses are in a publicly accessible <u>repository</u>. The algorithm delivered trial-by-trial estimates of pause onset, pause offset, the weights of the evidence for the onset and for the offset. It also delivered the duration of the longest inter-spike interval between these estimates, which may be considered an estimate of the depth of a pause. It also delivered the latency from pause onset to the onset of the longest inter-spike interval. This may be considered a measure of the abruptness of pause onset. Finally, it delivered the interval from the estimate of pause onset to the estimate of pause offset, which is an estimate of the width of the pause.

## Experiment 1: Acquisition of the Conditional Pause

Recordings where made from ten Purkinje cells in the eyeblink-controlling microzone within the C3 zone of the ferret cerebellar cortex during a conditioning protocol in which the CS was stimulation of the mossy fibers at 50 Hz. The US was direct stimulation of the inferior olive (n=5) or climbing fibers (n=5) with two short bursts of 5 pulses each, delivered at 500 Hz, with an interval between the bursts of 12 ms (first 7 subjects) or 4m (last 3 subjects). The ferrets were decerebrated before the experiment began.

In this and all subsequent experiments, each Purkinje cell was identified as an eyeblink cell by the presence of microzone-defining short-latency (10-12 ms) complex spike responses to brief electrical stimulation of the periocular receptive field (Hesslow 1994a&b)

In Cells/Subjects 1 through 7, CS stimulations terminated after 0.3s, and the US onset occurred .02 seconds later. In the last three Cells/Subjects, US onset occurred 0.2s after CS onset and the CS stimulation continued through and beyond US onset and offset. This was done to deconfound the effects of US onset from those of CS offset.

Recordings were initiated in the naive state, i.e., before the animal had been exposed to any (or only a few) paired stimulus presentations and continued for up to 4 hours, until a conditional pause was apparent over a sequence of trials. The inter-trial interval (the interval between CS onsets) was 15s.

## Results of the Analysis

### Complex and Variable Course of Conditioning

Figure 2 shows two raster plots with complete trial sequences. They are chosen to illustrate the complexity and diversity of what is seen during the training period, when the conditional pause develops

**Figure 2.**
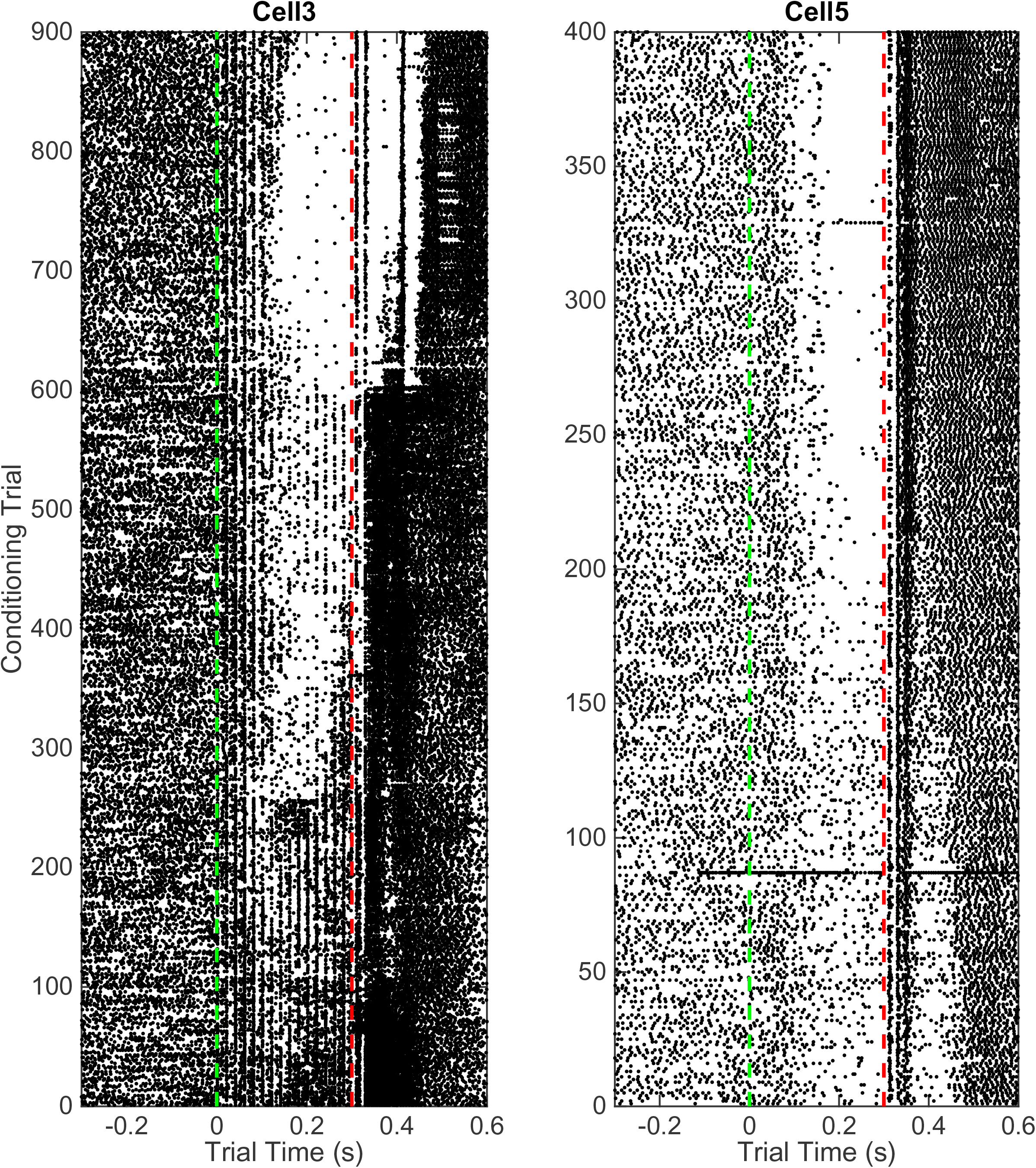
Raster plots. Each dot is a spike. Each horizontal line of dots is a trial. Vertical green dashed line indicates CS onset at trial time 0; vertical red dashed line indicates US onset.

In the left panel, one sees vertical lines of spikes at 20 ms intervals during the CS. These are clearly driven by the input, which was pulsatile mossy-fiber stimulation at 50 Hz. Early in training, the overall frequency of spikes is greater during the CS than before its onset. After about Trial 50, there are hints of a fall in frequency about half way through the CS, but this fall is more than offset by a clear increase in firing frequency in the first 100 ms. After 250 trials, a pause is evident, and the spikes elicited by the stimulation pulses begin to be suppressed during this pause. After 600 trials, this suppression is almost complete. In the right panel, there is no evidence of spikes elicited by the stimulation pulses. The pause is evident from the beginning, but it becomes more pronounced after about 180 trials.

The responses to the two brief bursts of US stimulation of the climbing fiber differ markedly between the two panels, and these responses to the US evolve in complex and disparate ways over the course of training. In some records, pronounced variations in firing rate are seen for several hundred milliseconds after US stimulation. These complexly evolving post-US variations in firing differ markedly from cell to cell. We do not attempt to analyze them in the present work.

### Non-Stationarity in Basal Firing Rate

There is substantial trial-to-trial variability in the firing rate prior to CS onset. Computation of the Fano factors for the spike count in the 1 s window immediately preceding CS onset shows that this variability is not consistent with a stationary Poisson process (Figure 3a). If it were, the Fano factor, which is the ratio of the variance in the spike counts to the mean count, would be close to 1, but in fact the distribution of Fano factors lies well beyond the limits expected from a stationary Poisson process (Figure 3a).

**Figure 3.**
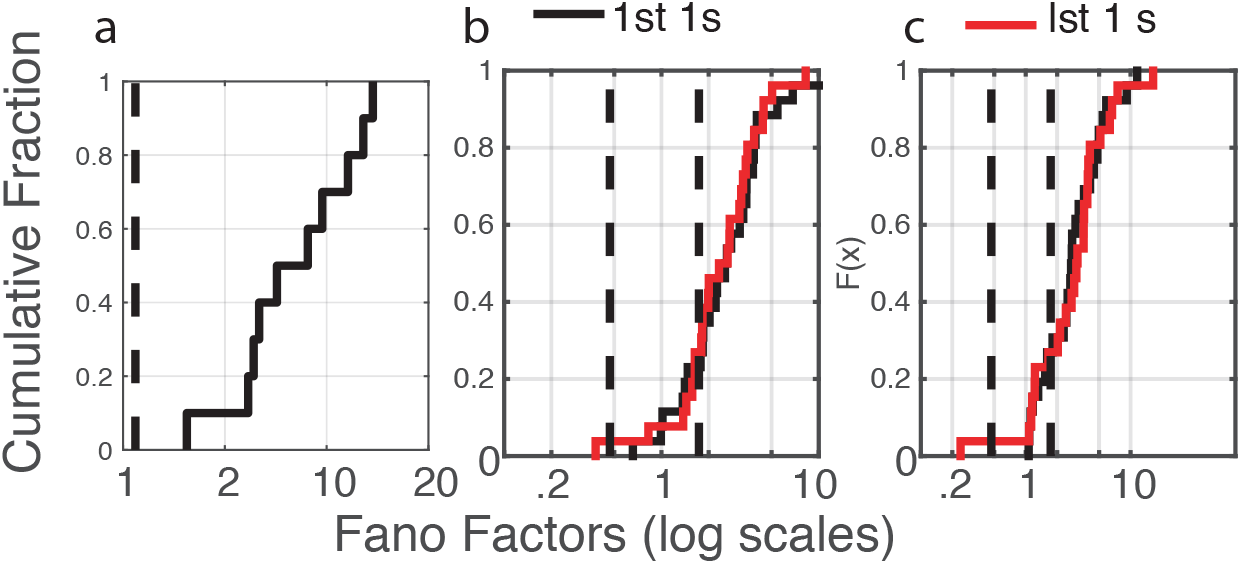
The cumulative distributions of Fano Factors reveal the marked between-trial non-stationarity of Purkinje cell firing patterns. The Fano factor is the ratio of the variance in the counts to their mean. In a stationary Poisson process, this statistic will be close to 1. The dashed vertical lines indicate an alpha of .01. Given the observed firing rates and sample sizes (numbers of trials), the Fano factor has a .01 probability of exceeding this limit if the process is stationary and Poisson (Eden & Kramer, 2010). a. The cumulative distribution of the between-trial Fano factors for the 10 cells in Experiment 1. The count window was the 1s immediately preceding CS onset. b & c . From Pre- (b) and Post- (c) CS spike trains in Experiment 2. The distributions in black are from the 1^st^ 1s; those in red from the last 1s of pre- and post-US spike trains lasting longer than 2s.

The evidence in Figure 3 of a basal firing rate that fluctuates from trial to trial raises the question of the time scale of this non-stationarity. We repeated the Fano Factor calculations using spike counts from successive 0.1 s windows in the pre-trial period of each trial—an order of magnitude smaller time scale than that in Figure 3. Because spike recording was turned on at varying intervals prior to CS onset, the number of such windows varied from 15 to 40. The bulk of these distributions fell within the plausible limits for a stationary Poisson process, although in several cells a substantial portion of the Fano factors were sub-Poisson (indicating less variance in the spike counts than would be the case if they were generated by a Poisson process). The distribution of interspike intervals is, however, not exponential; they have a much fatter tail.

### Variable Course of Acquisition

The common impression that behaviorally measured conditional responses develop gradually is an artifact of averaging across trials and subjects. The conditional response in most conditioning paradigms—both Pavlovian and instrumental—appears abruptly in most subjects (Gallistel et al., 2004; Papachristos & Gallistel, 2006). Its appearance is best visualized by means of a cumulative record of the trial-by-trial differences between the rate of responding prior to CS onset and the rate of responding during the CS (Figure 4). The slope of a cumulative record of a sequentially observed variable is the average value of that variable at a given point in its evolution. Early in conditioning—in the naive subject—the slope of the cumulative differences is usually 0 or even negative (because some subjects become wary in the presence of a novel stimulus). When the conditional response appears, the slope of the cumulative record becomes positive.

**Figure 4.**
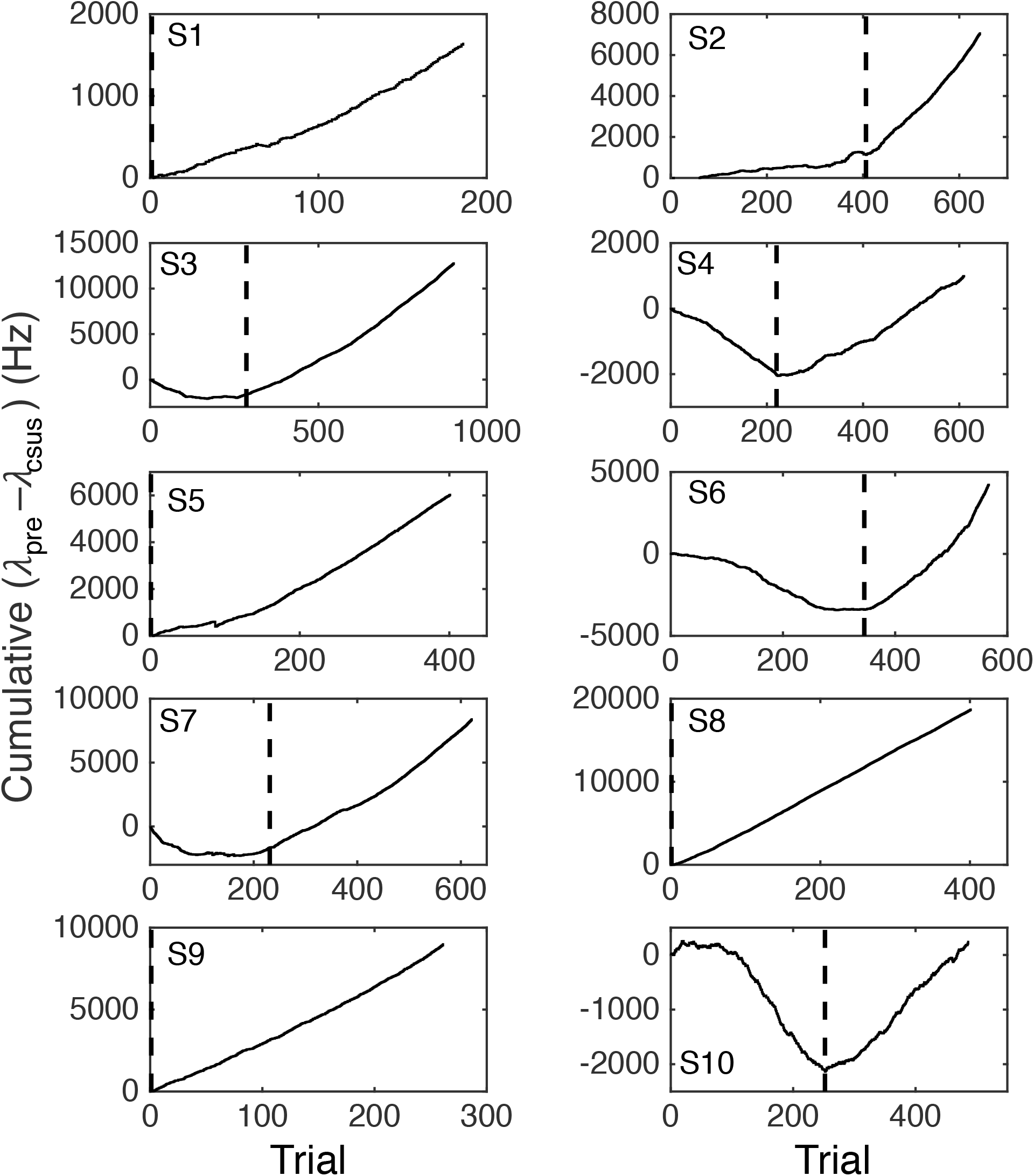
Cumulative records of the difference in firing rate between the immediately pre-CS interval equal in duration to the CS-US interval and the CS-US interval. An upward slope occurs when the average firing rate during the CS-US interval is less than during the interval preceding CS onset. Dashed vertical lines indicate the trial beyond which this average difference exceeded 10 Hz. This point is an estimate of the trial at which the conditional pause appeared (trials to acquisition).

The advantage of visualizing acquisition by means of a cumulative record is that there is no averaging. Hence, there is no smoothing; the more abrupt the change in behavior, the more abrupt the change in the slope. There are well-established algorithms for objectively identifying changes in slope (Gallistel et al., 2004). We can visualize the emergence of the conditional pause in the firing of a Purkinje cell by making a cumulative record of the difference between the pre-CS firing rate and the firing rate during the CS. And, we can apply to these records, the algorithm for identifying the changes in the slope (Figure 5).

**Figure 5.**
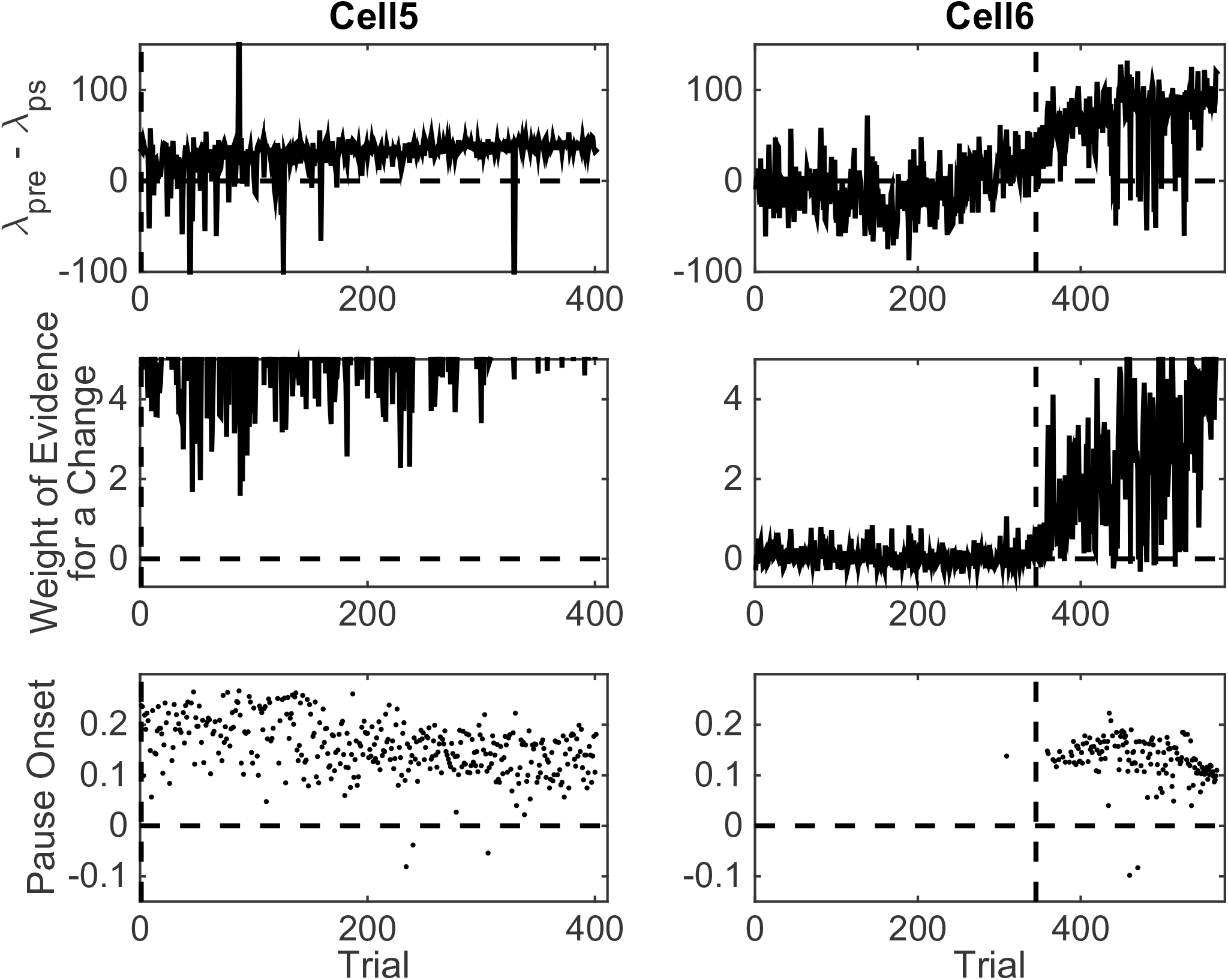
Pause statistics for two cells as a function of training trial. Top row: the difference between the pre-CS firing rate and the firing rate during the CS. Middle row: weight of the evidence log_10_(odds) for a change in firing rate between the pre-CS interval and the CS interval. Bottom row: Estimated pause onset latency (s).

Peristimulus histograms with 10 ms bin widths (see Figure 1b) enable the computation of the momentary probability of a spike within each bin. From such histograms we estimated three quantities that enter into the computation of an estimate of the pause onset time on a trial by trial basis: 1) the 10 ms wide interval at which the momentary probability of a spike is minimal (the bin with the lowest bar, see Figure 1b); 2) the momentary spike probability at that low point (the height of the lowest bar); 3) the momentary spike probability over the pre-CS interval (the average heights of the bars to the left of the 0 at CS onset, see Figure 1b).

Because the offset of pauses during training tends to coincide with the onset of US stimulation, pause offset cannot be estimated on a training trial; only pause onset can be. The Bayesian algorithm for estimating pause onsets operates on a binarized version of the spike train. Trial time is divided into successive 1ms “moments.” The 1ms width of these moments is chosen to be so narrow that at most 1 spike can occur during any one moment. Binning trial time in this way yields a binary string in which there is a 0 in every moment that did not contain a spike and a 1 in every moment that did. Binarization converts firing rates to momentary spike probabilities: the higher the firing rate, the greater the momentary probability of a spike. Then, the problem of estimating pause onset becomes one of estimating where the momentary probability of a spike increases as one looks back from the time where that probability is minimal (the retrospective binary sequence).

The algorithm computes the relative likelihood of two stochastic models for the retrospective binary sequence, a model in which the momentary probability is constant, and a model in which there is a step increase in the momentary spike probability as one looks back through the spike train from the low point within the CS. In the course of computing the second model, the algorithm determines the maximally likely location of this step and the strength of the evidence that it in fact exists. The evidence for or against its existence is the log of the odds in favor of the 1-change model as opposed to the no-change model (the null hypothesis in change detection). A log odds of 1 corresponds to odds of 10:1 in favor of the change model; a log odds of −1 corresponds to odds of 10:1 in favor of the no-change model. The log of the odds is called the weight of the evidence; weights of 2 and −2 correspond to 100:1 odds; −3 and 3, to 1,000:1 odds and so on. For the details of the algorithm, see the Appendix.

Figure 5 graphs pause acquisition statistics for two of the 10 cells. The dashed vertical lines are at the same locations in these plots as in Figure 5, namely, at the estimated trial of acquisition. Cell 5, whose statistics are plotted in the left column, was one of the 4 cells whose pause-acquisition was estimated in Figure 5 to occur at Trial 2, that is, after a single experience of the CS-US interval. Consistent with this, we see in the top left panel of Figure 5 that in this cell on the great majority of trials, *λ*_pre_, the firing rate during the pre-CS interval was higher than *λ*_ps_, the firing rate during the pause. The sign of the difference in firing rate was reversed in several early trials, but as training progressed, the trial-to-trial variability in the firing-rate difference decreased markedly. We see in the left middle panel that the evidence for this change in the firing rate was very strong even at the beginning, although it clearly became stronger still as training progressed and the pause became broader. The upper limit on the weight of the evidence (the ordinates of the middle panels) has been set to 5, which corresponds to odds of 100,000:1. The plot rises above this limit on most later trials. Even on the earliest trials, it is not infrequently above this limit. Finally, we see in the bottom left panel that the central tendency of the pause onset latency became shorter as training progressed and the variability in this onset latency decreased.

Cell 6, whose statistics are plotted on the right-hand column in Figure 5 was one of the 6 cells with an estimated pause-acquisition trial greater than 2. In the top-right panel, we see that on trials prior to this estimate, the *λ*_pre_– *λ*_ps_ difference fluctuated around 0. In the middle-right panel, we see that the evidence for a change in firing rate was weak and often negative (that is, the odds favored the model in which *λ*_pre_– *λ*_ps_ = 0). For a pause onset to be detected on a given trial, the *λ*_pre_– *λ*_ps_ difference must be positive and the evidence of a change must exceed 1 (10:1 odds) in favor of a step decrease in firing rate at the estimated location). In the light of the data in the top two panels on the right, we are not surprised to see that, with one exception, pause onsets were not detected until after the dashed vertical. Thus, what we see in these plots is consistent with the estimated trials on which a conditional pause was acquired (dashed verticals in Figure 5), and this is true for the other six cells as well.

Two aspects of the plots in Figure 5 may seem puzzling: First, some pause onsets are negative. Second, there is sometimes strong evidence of a change on a given trial but no pause onset is detected. Pause onsets can be negative because the distribution of inter-spike intervals in the spontaneous firing of Purkinje cells has an extremely long tail. Although the modal inter-spike interval is often less than 10ms, inter-spike intervals longer than 100ms occur with some frequency. The pause-onset latency in a cell trained with a 300ms CS-US interval hovers near 100ms (see Pause On Lat panels in Figure 6). When CS onset occurs soon after the beginning of a spontaneous inter-spike interval greater than 100ms, the shut-down in the firing produced by the CS occurs during that interval. In that case, the onset of the pause will appear to precede the onset of the CS.

**Figure 6.**
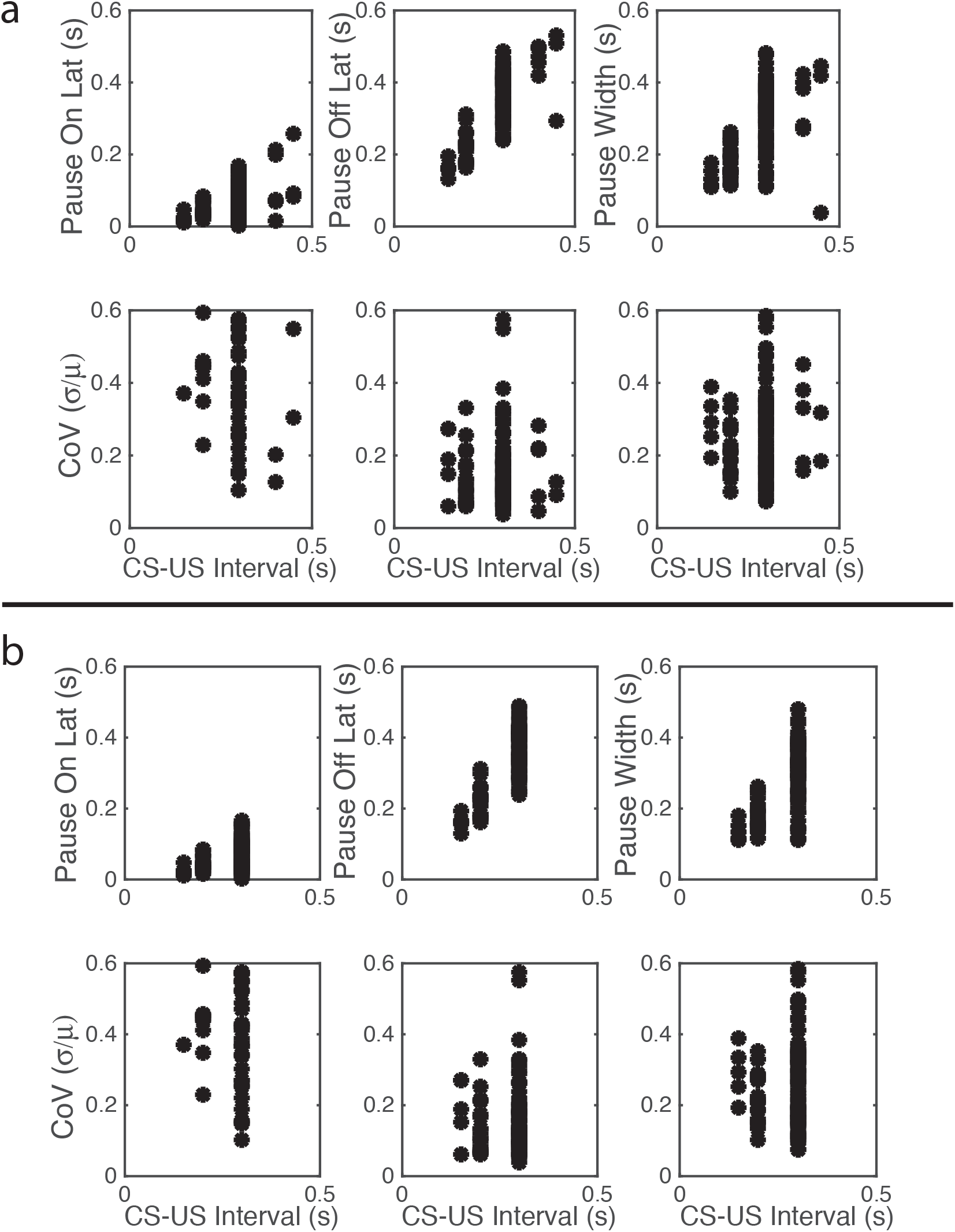
Scatter plots of mean pause-onset latencies, mean pause-offset latencies and mean pause widths as a function of the CS-US training interval and their coefficients of variation (CoV). a. Experiment 2, in which the CS was stimulation of the dorsum of the paw. b. Experiment 3, in which the CS was direct stimulation of the parallel fibers, immediately presynaptic to the Purkinje cell. Note the close similarity of the results in a and b, despite the radically different way in which the signal arriving at the Purkinje cell was generated.

A pause onset is not detected despite strong evidence of a change in ps when the algorithm encounters a clear negative step in the *λ*_pre_– *λ*_ps_ difference when looking backward in the spike train. The computation of the strength of the evidence for a change is oblivious to the sign of the change, but a pause onset is detected only on trials where *λ*_pre_– *λ*_ps_ >0. On some rare anomalous trials, the firing rate increases during the CS rather than decreasing. On those trials, a pause is not detected.

## Discussion

These cellular-level data on trial to acquisition and pause onset latency are broadly consistent with behavioral level data. We postpone the discussion of the data on onset latencies until after the analysis of the next data set, which is much larger, and has a much wider variation in the CS-US interval. Here, we discuss only trials to acquisition.

Trials-to-acquisition vary greatly between subjects in eye blink conditioning, as in most other forms of conditioning (Gallistel et al., 2004; Papachristos & Gallistel, 2006). In the conditioning of these cells, the inter-trial interval was very short (15s). Trials to acquisition are generally inversely proportional to the inter-trial interval; with intervals this short, the number of trials required often runs into the hundreds (Gallistel & Gibbon, 2001; Gibbon & Balsam, 1981).

It may seem remarkable that a pause often appears after a number of trials so small that the number of trials to acquisition cannot be estimated by a change-detection algorithm, in which case an algorithm for estimating trials to acquisition will set the number at 0. 0 trials to acquisition is analytically impossible. There must be at least one trial before a response based on information communicated only in a trial can be observed. In associative learning protocols that use foot shock, one trial learning is common. In eye blink conditioning, however, the lower end of the trials-to-acquisition range is around 10 trials, at least with non-human subjects. Several considerations are relevant in considering what conclusions to draw from the fact that in some preparations there is evidence of a conditional pause after the first training trial. The most important of these considerations is conceptual; it has to do with the difference between a plastic conception of memory-formation in associative learning and an inscriptional conception.

The importance of the Purkinje cell pause-conditioning phenomenon is that it provides neuroscience with an example in which a specifiable quantitative fact gleaned from experience has been localized to the cellular level. It appears necessary to think of this phenomenon in inscriptional terms; the conditioning protocol has inscribed the duration of the CS-US interval into a medium intrinsic to the Purkinje cell (the engram). This conceptual framework differs fundamentally from the framework in which the phenomena of learning and memory are treated in the neurobiological literature, for reasons we now pause to explain.

The neurobiology of learning and memory is focused on synaptic plasticity (Martin & Morris, 2002; Poo et al., 2016). That focus reflects a commitment, witting or unwitting, to a behaviorist conception of learning and memory. In this conception, experience does not inscribe facts; it molds circuits. It alters the brain’s wiring by changing synaptic conductances. The changes in the synapses alter signal flow in such a way as to change the brain’s input-output function; they do not encode a fact extracted from experience. Thus, in work focused on mechanisms of synaptic plasticity (Poo et al., 2016, for example), there is no attempt to say how the mechanisms considered could encode a quantitative fact (Gallistel & Matzel, 2013; Gallistel, 2017; Langille & Gallistel, 2020).

The coding question is unavoidable in an inscriptional theory of learning and memory, because the facts (about, for example, interval durations) must be inscribed and read in accord with some code, just as the program for building an organism is inscribed in its DNA in accord with a code and read from that DNA by code-specific molecular machinery. When thinking in inscriptional terms, the notion of memory “strength” makes little sense. A fact is either legibly inscribed or it is not. If the first experience of a CS-US interval does not legibly inscribe the duration of that interval, then there is no way the brain can know that a second experience of the same interval is in fact the same as the first experience. As in anterograde amnesias, every experience of the same duration, no matter how often repeated, is a novel experience of that duration (Kritchevsky, Squire, & Zouzounis, 1988; Milner, Corkin, & Teuber, 1968; Siegert & Warrington, 1996). This consideration seems to require the assumption that in all conditioning protocols in which the subject learns that the CS-US interval has a fixed value, single experiences of that interval inscribe in legible form the duration of the CS-US interval. If the first experience of that interval did not legibly inscribe its duration, then every subsequent experience of the same interval would be no different from the first experience. There would, therefore, be no way for evidence to accumulate that the CS-US interval was constant from trial to trial; hence predictable. Nor would there be any way for the brain to distinguish trial-to-trial variability in its measurements of a fixed CS-US interval from actual variations in the interval itself, in those protocols where the CS-US interval varies. Rodents do, however, make this distinction (Kheifets, Freestone, & Gallistel, 2017; Li & Dudman, 2013).

Given this consideration, we are not surprised that evidence of having committed the duration of the CS-US interval to memory is apparent in some cells after a single experience of that interval. In all 10 cells, there was evidence for a further evolution over many trials of its response to the CS (see, for example, the bottom left panel in Figure 5). We assume that it is these further evolutions and the emergence of a conditional response in more than one Purkinje cell that explains the emergence of a behaviorally observable response.

## Experiment 2: CS is Stimulation of the Dorsum of the Paw

The second data set comes from 106 cells recorded from 54 decerebrated ferrets. The CS in the conditioning protocol was pulsatile stimulation at 50 Hz through an electrode on the dorsum of a forepaw. The US was stimulation of a climbing fiber with two 5-pulse bursts of pulses at 500 Hz, with a 10 ms interval between the two bursts. The CS stimulation terminated at US onset. The CS-US interval varied between subjects from as short as 0.15s to as long as 0.45s. The interval from CS onset to CS onset varied from as short as 6s for some subject to as long as 17 seconds in others. When a clear pause was observed, delivery of the US was discontinued, and 20 successive probe trials were run with CS alone to obtain the data on which the analyses here reported are based.

In about half the subjects, the electrode was then advanced to find a nearby Purkinje cell that had also been conditioned, and 20 further probe trials were run while recording from that cell. In some cases, as many as 6 trained cells were recorded in a single subject.

## Results of the Analysis

We again computed the Fano Factors for the spike counts in two pre-CS and two post-CS windows of 1s width, for all those cells in which the pre-CS and post-CS spike trains were both recorded for more than 2s. The two windows were the first 1s and the last 1s of such spike trains. The cumulative distributions of the Fano factors are shown in Figure 3 b&c. The pre-CS and the post-CS distributions are the same (compare b & c in Figure 3), as are the distributions for the first and last 1s windows (compare black and red distributions within panels).

Seventy-five percent of the Fano factors are beyond the upper limit on a plausible Fano factor from a stationary Poisson process. Moreover, and perhaps more importantly, even for the rare cells for which all 4 Fano factors were within the plausible Poisson limits, the distributions of interspike intervals are not well fit by the exponential distribution that describes intervals generated by a Poisson process. Like the distributions for all the other cells, they have a prolonged fat tail.

In sum, the endogenous process that generates the high rate of spontaneous spiking in Purkinje cells is not a Poisson process and it is not stationary. While the bulk of the interspike intervals fall in a range of 5-15 ms, intervals of 40-150 ms (and occasionally even much longer) are reliably present in samples lasting more than a few seconds. Given the importance of this cell with its very unusual firing pattern and its very unusual complex spikes that probably play a central role in learning, intensive study of the mechanism that generates this firing may pay big dividends. The unusual size of the cell should facilitate such studies.

### Pause Statistics

Using our algorithm for finding pause onsets and offsets (see Appendix), we found trial-by-trial pause onsets and offsets for the data sets from each of 106 cells. From the offsets and onsets, we computed the pause widths. We also computed the maximum interspike interval between each pause on and pause off and the latency from pause on to the onset of the longest interspike interval.

The top row of Figure 6a gives scatter plots of the pause-onset latencies, the pause-offset latencies and the pause widths (the difference between the two latencies), while the bottom row gives the coefficient of variation (CoV) in these statistics. The CoV is the ratio between the standard deviation of a random variable and its mean. In time-scale-invariant measures, measures that obey Weber’s Law, the standard deviation in the measure increases in proportion with the mean, so the CoV is constant. We do not graph the fifth basic statistic—the latency from pause onset to the beginning of the longest within-pause inter-spike interval—because most pauses begin with the longest within-pause inter-spike interval, regardless of the duration of the CS-US interval. From this and other aspects of the data, we conclude that the unusually long interspike interval *is* the pause.

### Pause Statistics Scale with the Training Interval

The most conspicuous aspect of the results in Figure 6a is that the pause onset and pause offset and the interval between them become progressively longer as the CS-US interval during training increases. A second feature is that CoVs for these intervals are constant. In other words, these statistics exhibit scalar variability. Scalar variability is ubiquitous in behavioral timing (Gallistel & Gibbon, 2000; Gibbon, 1977, 1992; Gibbon, Church, & Meck, 1984; Simen, Rivest, Ludvig, & Killeen, 2013; White et al., 2000). It is a manifestation of Weber’s Law and of time-scale invariance, both of which are quantitatively important aspects of associative learning (Gallistel & Gibbon, 2000). The range covered by the CoVs of the pause offset latencies is the same as that observed in behavioral experiments on the timing of conditional responses (Gallistel & Gibbon, 2000).

### Pause Onset Latency Can Be Extremely Short

The onset latencies are remarkably short when cells are trained with a 0.15s CS-US latency. This latency is close to the shortest CS-US interval that will produce a conditional eye blink (0.1s), which is also the shortest interval that will produce a conditional pause in the Purkinje cell (Jirenhed & Hesslow, 2016; Salafia, Mis, Terry, Bartosiak, & Daston, 1973; Schneiderman & Gormezano, 1964). The median of the median onset latencies in the group trained with a 0.15s CS-US interval is slightly less than 20 ms. Thus, in 50% of the cells, half the pauses began before the delivery of the second pulse in the train of CS pulses that was delivered to the dorsum of the forepaw. In the much more numerous group of cells trained with a 0.2s CS-US interval, the median of the median pause onset latency was 40ms and the median of the 1^st^ quartiles was just over 20 ms. Thus, on half the trials in this condition, the pause began at or before the delivery of the 3^rd^ pulse in the train of CS pulses, and on slightly less than 25% of the trials it began at or before the delivery of the 2^nd^ pulse in the CS stimulation of the forepaw.

### Pause Onsets and Offsets Are Step-Like

The first inter-spike interval within the pause is more often than not the longest inter-spike interval within the pause. This is a major reason for our conclusion that on any given trial a single unusually long interspike interval *is* the pause. This hypothesis explains why the distributions of pause widths and of the longest inter-spike intervals within the pause are so similar, as may be seen in Figure 7, which plots the cumulative distributions of the pause-width-distribution quartiles (left columns) and the cumulative distributions of the quartiles of the longest-within-pause inter-spike interval distributions (right column). The onset latency and duration of this unusually long inter-spike interval are determined by the memory of the previously experienced CS-US intervals (the engram).

**Figure 7.**
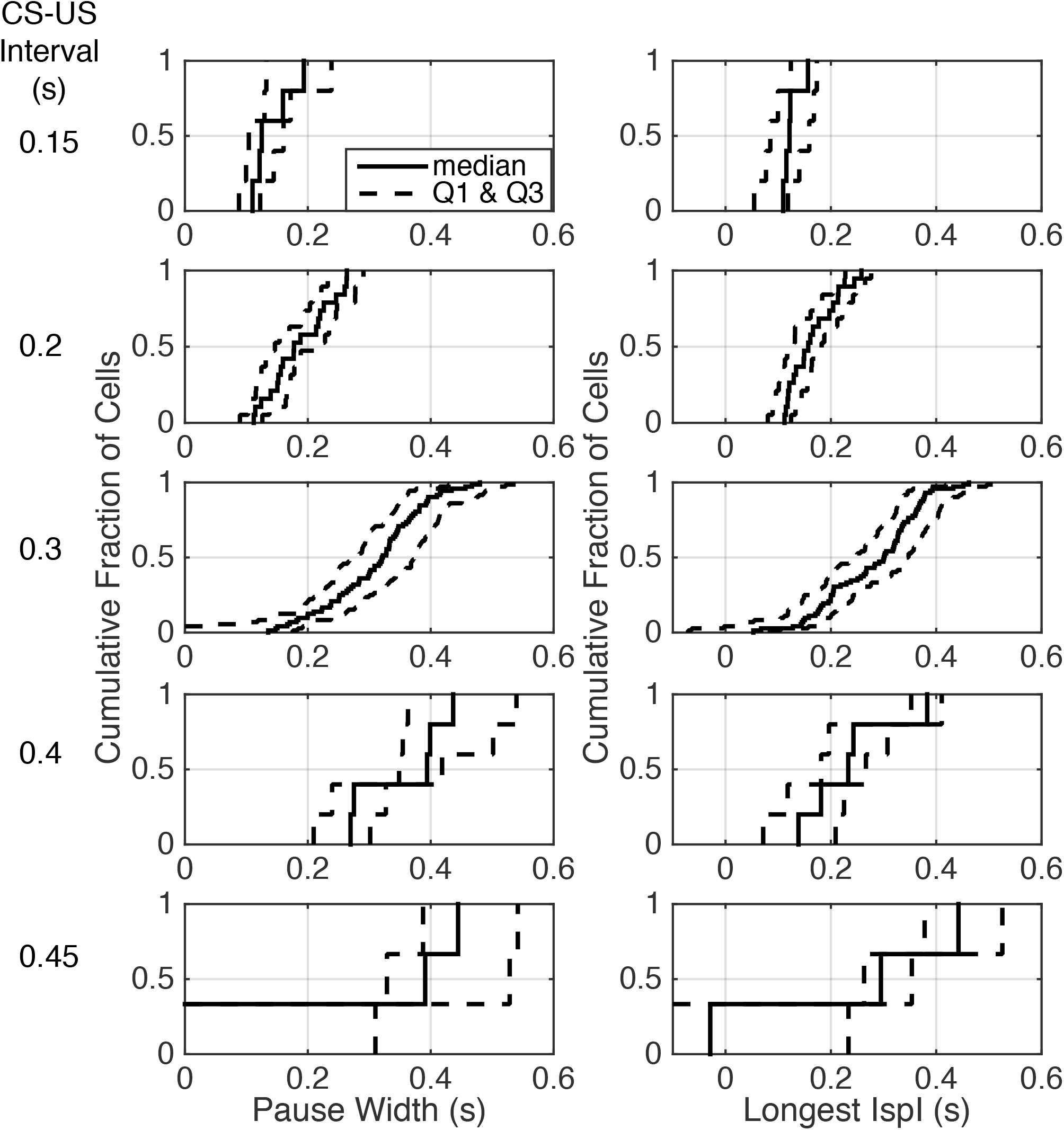
Quartiles of the within-group distributions of pause width (left columns) and longest within-pause inter-spike interval (right column). Cells are grouped by the CS-US interval during their conditioning (given to left of each row). The second quartile (the median) is plotted with a solid line; the first and last quartiles with dashed lines. The strong similarity between the panels on the two sides of this figure is evidence that a single unusually long interspike interval constitutes the conditional pause.

To check whether there was evidence of any lengthening of the inter-spike intervals after CS onset but before what our algorithm identified as the onset of the pause, Figure 8 gives scatter plots for the successive inter-spike intervals encountered when looking backward from pause onset to CS onset in a representative sample of cells trained with a 0.3s CS-US interval. These plots do not suggest any reliable upward trend in the inter-spike intervals between CS onset and pause onset. Figure SF1 in the Supplementary Material gives a similar plot for the data from a 0.4s CS-US interval; again, there is no evidence of an increase in the interspike intervals that precede the unusually long interval.

**Figure 8.**
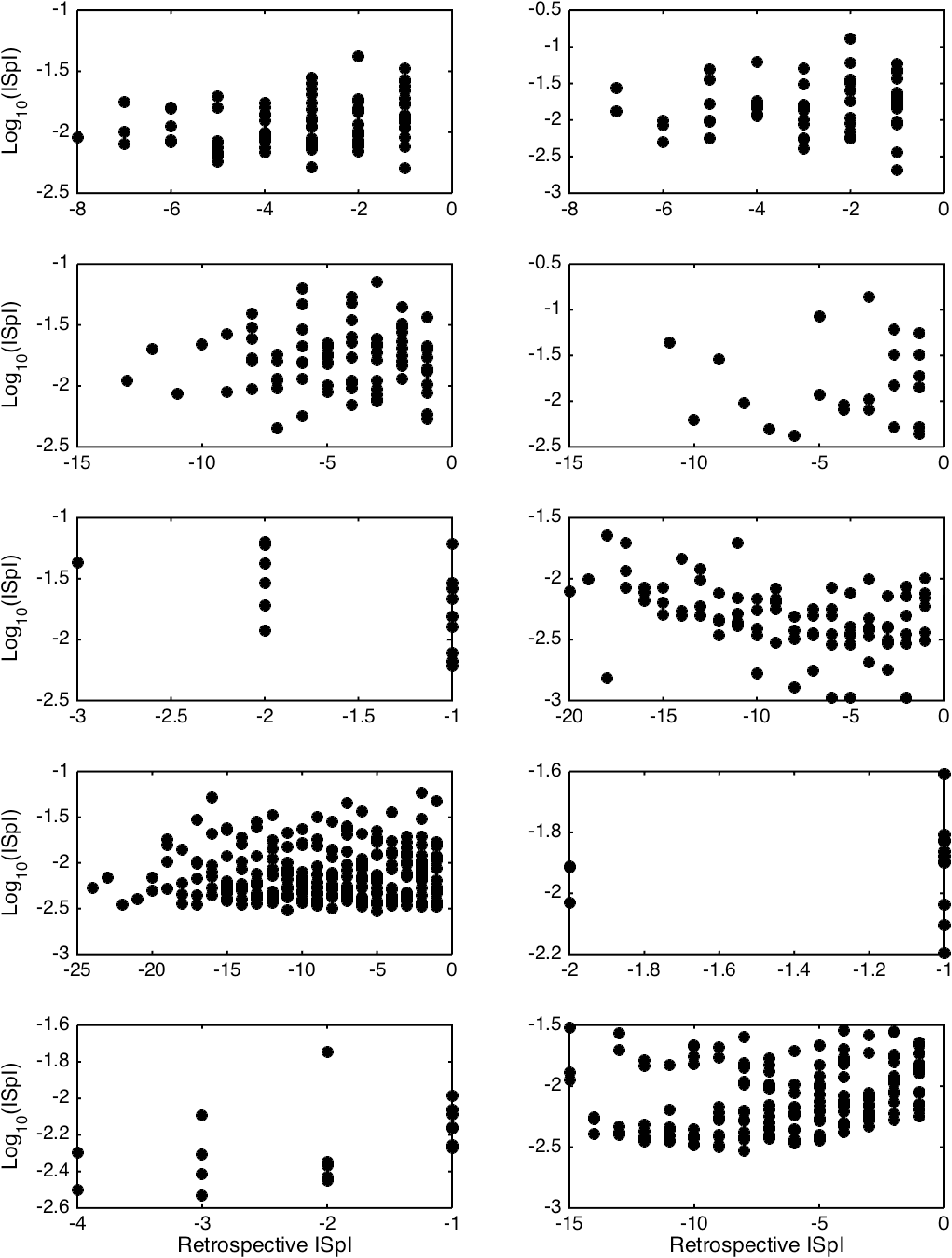
Successive inter-spike intervals encountered when looking backward from pause onset to CS onset. Pause onset is at 0 at the right edge of each plot; –1 is the first interval encountered; –2, the second, and so on, as one progresses leftward (backward in time from pause onset). Each plot is for a cell trained with a 0.3 CS-US interval, because pause onset was generally well after CS onset in this condition. If there were a gradual increase in the inter-spike interval prior to the onset of the longest inter-spike interval, these scatter plots would drift upward, but they do not.

As a further check on whether the interspike intervals tend to grow longer as the appearance of the unusually long interspike interval draws nearer, we computed the linear regression of the interspike intervals backward from the onset of the longest interspike interval within the pause to the onset of the CS. We did this for all those trials in the CS-US >=0.3s conditions in which there were at least 5 interspike intervals between CS onset and the estimated onset of the pause. The bulk of the slopes in these regressions are positive, that is, the interspike intervals farther back in the retrospective sequence (closer to CS onset and farther from the onset of the unusually long interspike interval) tended, if anything, to be *longer* than those that are earlier in the retrospective sequence (closer to the pause onset). The tendency is, however, very weak, because in 75% of the regressions, the lower confidence limit on the slope is negative—thus, the confidence interval includes 0—and 80% of the regressions explain less than 7% of the variance. In sum, there is no tendency for the inter-spike interval to grow longer as the onset of the pause draws nearer; the very weak and unreliable tendency that exists is in the opposite direction.

We conclude that the conditional pause in the firing of the cerebellar Purkinje cell consists more often than not of a single unusually long inter-spike interval whose onset and offset latencies are a scalar function of the CS-US training interval. There is no sign of the graded increase in inter-spike intervals that would be expected if a gradually strengthening inhibitory synaptic input increased postsynaptic membrane polarization in a temporally graded manner.

### Correlations Among Pause Parameters

Figure 9a shows for each CS-US interval group the pairwise correlations among the 4 pause parameters: pause onset, pause offset, pause width and the longest within-pause inter-spike interval. The structure is the same regardless of the CS-US training interval (compare across panels in Figure 9a). The correlation between pause onset latency (↑) and pause offset latency (↓) is highly variable between cells within a group, but the central tendency is close to 0. By contrast, the correlation between pause width (W) and pause onset latency is consistently negative, often strongly so: a late onset predicts a short pause. Because the pause very often begins and ends with the beginning and end of the longest within-pause inter-spike interval (M), it is not surprising that a late onset also predicts that the longest inter-spike interval within the pause will be relatively short. In this same light, it is also not surprisingly that pause width and the longest within-pause inter-spike interval are strongly and positively correlated with pause-offset latency, and very strongly correlated with one another. We defer to the General Discussion a discussion of the implications of these correlations.

**Figure 9.**
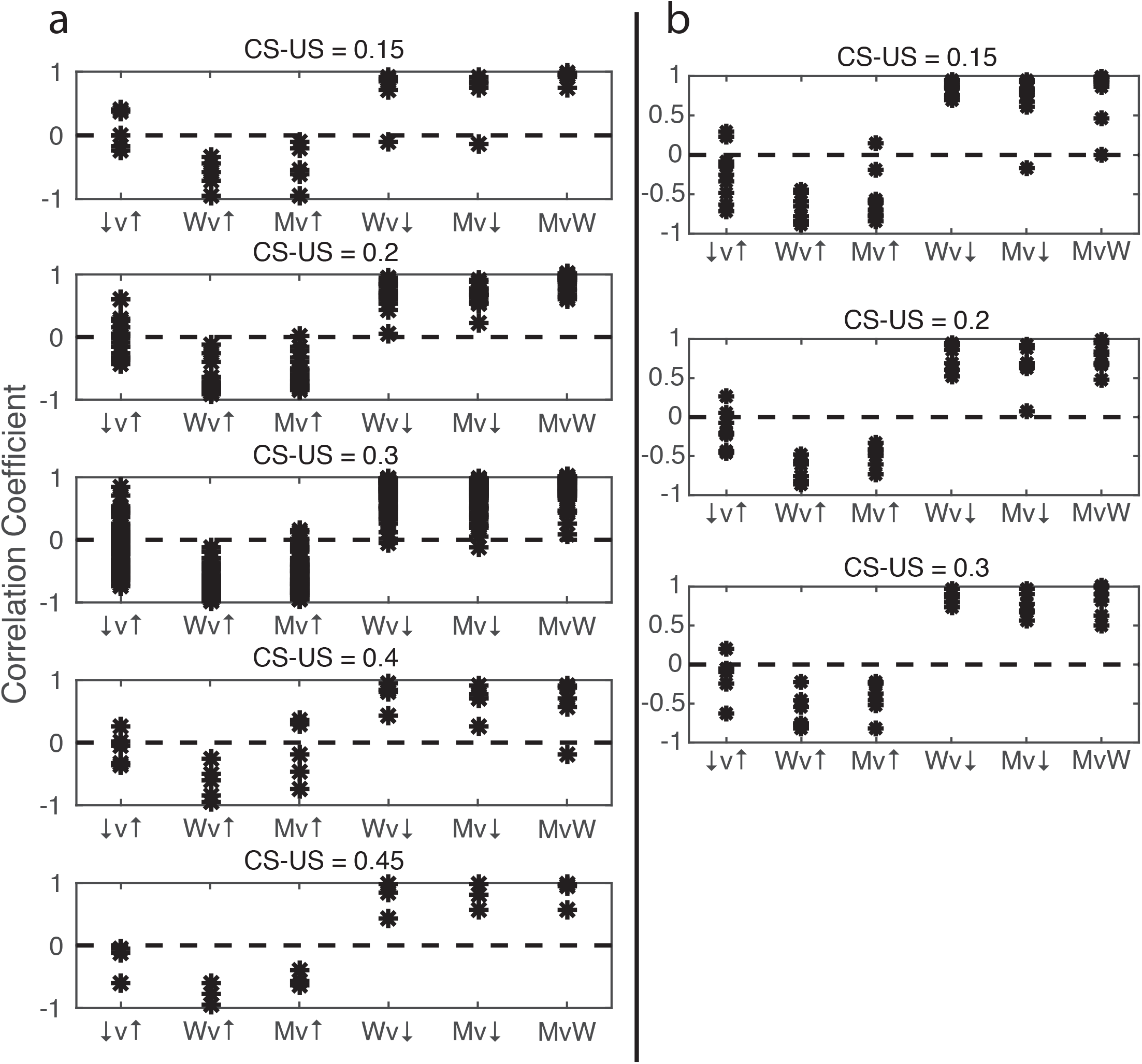
The pairwise correlation structure among four pause parameters: pause onset latency (↑), pause offset latency (↓), pause width (W), and longest within-pause inter-spike interval (M). The CS-US training interval varies between panels. a. Data from Experiment 2. b. Data from Experiment 3. Note the close similarity of the results in a and b, despite the radically different way in which the signal arriving at the Purkinje cell was generated.

## Experiment 3: CS is Stimulation of the Parallel Fibers

In this experiment, the CS was direct pulsatile electrical stimulation of the parallel fiber input to the Purkinje cell at 50 or 100Hz (0.1ms pulse width), and the US was, as usual, two very short bursts of very high frequency (500Hz) stimulation of the climbing fiber. In this experiment, there was direct control of the presynaptic signal to the Purkinje cell. In previous experiments, the CS signal was generated by stimulation delivered at a remove of two or more synapses from the Purkinje cell. Hence, it was possible that neurons intervening between the site of CS stimulation and the Purkinje cell provided a presynaptic signal with a rise and fall that might explain the pause. In this experiment, the presynaptic signal, the signal seen at the synapses between the parallel fibers and the Purkinje cell was under direct experimental control.

We here analyze 22 cells from this experiment. Nine of them were conditioned with a CS-US interval of 0.15s but a CS duration of 0.3s. In other words, CS offset was not coincident with US onset during training. More importantly, in the 20 probe trials with no US, which followed the conditioning, and which provide the data we here analyze, CS offset was not coincident with the time at which a US was anticipated but failed to occur. Thus, the termination of CS stimulation did not play a role in the generation of the pause offsets—the recovery of spontaneous firing at the time when a US was anticipated. This recovery occurred well before the termination of the CS stimulation.

Five of the 22 cells were conditioned with CS-US intervals of 0.2s but CS durations of 0.8s. Again, in these 5 cells, the termination of CS stimulation occurred long after the time at which a US was anticipated. Two more cells were conditioned with the same CS-US interval (0.2s) and with CS termination at US onset.

Finally, six cells were conditioned with CS-US intervals of 0.3s and US onset at CS termination.

## Results

The most striking thing about the results is their similarity to the results obtained with CS stimulation delivered to the dorsum of the paw (see Figures 6 and 9). The similarities are immediately apparent in the raw data, that is, in the raster plots like those in Figure 1 (<u>link</u>).

The general conclusion from this experiment is that when the CS signal arriving at the Purkinje cell is produced by direct pulsatile electrical stimulation of the immediately presynaptic afferents to the Purkinje cell, the quantitative characteristics of the pause are the same as when the pause is elicited by a sensory CS (stimulation of the dorsum of the paw). The pause is the same regardless whether the stimulation of the parallel fibers terminates at the time when the US is expected or continues for hundreds of milliseconds after that time. Finally, the pause is the same during probe trials even when the stimulation parameters differ radically in both duration and pulse frequency from those used during training (when the US was presented). These results strongly suggest that mechanisms intrinsic to the Purkinje cell itself determine the quantitative properties of the learned pause in its firing triggered by the onset of parallel fiber input, because the pause is in every way the same under conditions where it is unlikely that there is any time-varying input to the Purkinje cell during a CS.

## General Discussion

### Agreement Between Behavioral Measurements and Cellular Measurements

When attempting to establish the material basis for a mechanism known only from its behavioral effects, it is essential to establish a quantitative correspondence between properties of that mechanism established by measurements based only on its behavioral effects and properties measured by “more direct” non-behavioral means that bring one closer to the presumed locus of the unknown mechanism. For example, when Du Bois-Reymond asserted that the “action current” (now called the action potential), which he had discovered in nerve, was the physical realization of the nerve impulse (Du Bois-Reymond, 1848), it was clear that, for his hypothesis to be true, the velocity of the action potential had to be the same as the velocity of the nerve impulse. At about the same time as Du Bois-Reymond did his electrophysiological work, Helmholtz, his friend and colleague in the laboratory of Johannes Müller, measured the velocity of the nerve impulse in frog motor nerve and human sensory nerve (Helmholtz, 1850, 1852), using the difference in reaction time method, which remains a staple of cognitive neuroscience (Luce, 1991). In 1871, a student of Du Bois-Reymond did the behavioral measurements and the electrophysiological measurements on the same preparation (the frog sciatic-gastrocnemius preparation) and found the velocities to be the same within the errors of measurement (Bernstein, 1871), as required by his advisor’s hypothesis.

In attempting to establish the material basis of memory, the same considerations apply (Gallistel, Shizgal, & Yeomans, 1981): the quantitative properties of a putative cellular level manifestation of the engram must align with the quantitative properties established by behavioral experiment. Electrophysiologically measured synaptic plasticity fails to satisfy this requirement: No measured property of Long Term Potentiation (LTP), Long Term Depotentiation (LTD) or Spike Timing Dependent Plasticity (STDP) corresponds to any behaviorally measured property of associative learning (Gallistel & Matzel, 2013).

The only electrophysiologically measurable phenomenon whose quantitative properties align with the behaviorally established properties of the corresponding associative learning phenomenon is the conditional pause in the spontaneous firing of the cerebellar Purkinje cells, the cells that have been shown to control the timing of the conditional eyeblink (Heiney et al., 2014; Johansson et al., 2016). Jirenhed and Hesslow (2016) review and document the following quantitative correspondences: The numbers of trials required for the acquisition of the conditional response (CR) fall, in both cases, within the same range, as do the numbers required for its extinction. In the cellular preparation as in the behavior of the intact subject, CS-US intervals shorter than 80-100 ms do not induce a CR. In both, reacquisition of the CR following its extinction occurs much more rapidly than the original acquisition. Preliminary results obtained by one of us (Johansson) suggest that lengthening the intertrial interval causes the CR to appear after fewer trials, as it does for the behavioral CR in the intact rabbit (Gallistel & Gibbon, 2000, their Figure 10). And, of course, the timing of the cellular CR depends on the CS-US interval in the same way as does the timing of the behavioral CR. Its onset latency, offset latency and duration vary in the same way with the CS-US interval, and the offset of the conditional pause occurs at approximately the latency at which the US is expected, which is the time at which lid or membrane closure attains its maximum.

**Figure 10.**
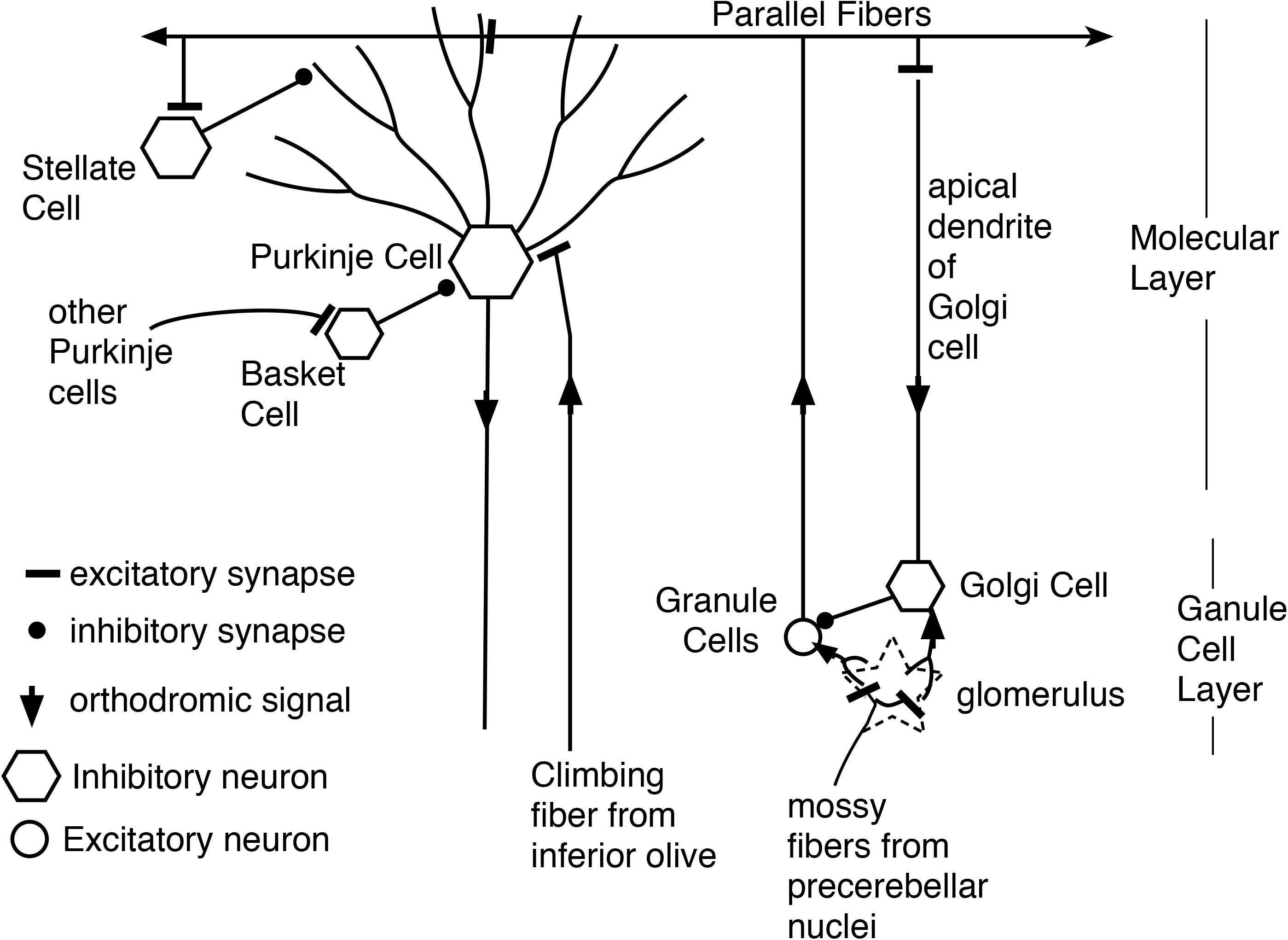
Cerebellar circuitry. Each parallel fiber contacts tens of thousands of Purkinje cells and each Purkinje cell receives input from hundreds of thousands of granule cells, but from only one climbing fiber. The parallel fibers also make excitatory synapses on the stellate cells and on the Golgi cells. The basket cells make powerful inhibitory synapses on the Purkinje cell soma; they get their excitatory input from other Purkinje cells. Not shown are the Lugaro cells, which are excited by Purkinje cell collaterals and which make inhibitory synapses on stellate cells, basket cells and Golgi cells.

The results we here report add to the list of quantitative correspondences. They show that the CoV for the offset latency in the conditional firing pause of Purkinje cells in the C3 microzone of the decerebrate ferret. Given that the cellular data come from decerebrate ferrets while the behavioral data come from intact rabbits, this may perhaps be regarded as a reasonable correspondence. It is, however, clearly desirable that both the cellular and the behavioral measurements be made on the same preparations. Also, of course, if the between-cell sources of variance are independent and if the behavioral variance depends on the pooling of the signals from *n* cells, then the behavioral standard deviation will be smaller than the cellular standard deviation by the square root of *n*.

The systematic increase in the central tendencies of the timing quantities and the proportionate increase in their variability are further examples of the many ways in which the quantitative properties of this cellular-level learning phenomenon match the quantitative properties that are known to obtain at the behavioral level (Jirenhed & Hesslow, 2016). This correspondence between quantities measured at the cellular level and corresponding quantities measured at the behavioral level stands in marked contrast to the situation with LTP (long term potentiation) and STDP (spike-timing-dependent plasticity). None of the quantitative properties of these cellular level phenomena agree with the behaviorally determined quantitative properties of associative learning (Gallistel & Matzel, 2013; Martin & Morris, 2002).

### Is There an Engram Intrinsic to the Purkinje Cell?

The engram for a CS-US interval is a hypothesized physical change at the circuit and/or the molecular level of a structure, which change is produced by the conditioning experience. The altered structure encodes the duration of the interval in a way that may be read on demand and may enter into computations with other remembered intervals. The hypothesized engram explains the well-established fact that the conditional response that emerges in Pavlovian and instrumental conditioning protocols is appropriately timed and the further fact that ratios and differences in protocol intervals are critical determinants of the conditioned behavior (Arcediano & Miller, 2002; Balsam et al., 2010; Barnet, Cole, & Miller, 1997; Denniston, Blaidsdell, & Miller, 1998; Gallistel & Gibbon, 2000; Kalmbach, Chun, Taylor, Gallistel, & Balsam, 2019; Matzel, Held, & Miller, 1988; Ward, Gallistel, & Balsam, 2013; Ward et al., 2012).

In most contemporary theorizing about the neurobiological mechanisms underlying the timing of conditional responses, there is no such engram. In lieu of an engram, most contemporary theories posit the selective formation of associations between response-generating neurons and an appropriate set of temporal grandmother neurons or grandmother circuits (Bareš et al., 2018; S. Grossberg & Schmajuk, 1991; Grossberg, 1991; Mauk & Buonomano, 2004; Meck, 2003; Yamaski & Tanaka, 2009). In their most extreme form—in timerless timing theories—even the temporal grandmother cells are dispensed with. Timerless timing theories posit only feed forward activities that culminate at different times because of the complex dynamics of neural circuits. The neurons at the end of the appropriate chain—the neurons or neuronal ensembles that become active at the right time—become selectively associated with the response-generating neurons (Hardy & Buonomano, 2018; Yamazaki & Tanaka, 2009).

The experiments in which the CS signal is produced by direct stimulation of the parallel fibers appear to rule out engramless models of timing. The crucial assumption in these experiments is that the parallel fiber input seen by the Purkinje cell directly reflects the train of evenly spaced stimulating pulses delivered to the parallel fibers. If that is the case, the dynamics of presynaptic circuits are irrelevant, because the experimenter has gained control of the relevant presynaptic input to the Purkinje cell. The signal that governs the timing of the pause arises within the Purkinje cell itself (Johansson, 2019). The parallel fiber elicits that signal by triggering the reading of the engram that encodes the interval.

That parallel fiber input evokes the conditional pause in the critical parallel-fiber-stimulation experiments is strongly implied by two pharmacological results cited in our introduction: 1) Blocking the mGluR7 receptor in the post-synaptic side of the parallel-fiber-to-Purkinje cell synapse blocks the pause. 2) Blocking the known inhibitory inputs to the Purkinje cell with a general-purpose GABA blocker (gabazine), given in a dose that blocks the very strong inhibitory effect of off-beam stimulation, does not block the elicitation of the conditional pause.

Also relevant is that, to our knowledge, neither the granule cells from which the parallel fibers originate, nor any other neurons in the cerebellum have been shown to exhibit the properties required of temporal grandmother cells.

Given the far-reaching implications for further research of the conclusion that there is an engram for the CS-US interval intrinsic to an intracellular biochemical cascade that begins with the activation of the mGluR7 receptor, it is important to consider whether the experimental facts so far obtained might permit of an alternative conclusion.

One such alternative might begin with the fact that direct stimulation of the parallel fibers triggers antidromic volleys of action potentials, as well as orthodromic volleys. Might these antidromic signals reach the Purkinje cell by some path other than the directly stimulated parallel fibers? Several colleagues, reviewers and speakers at scholarly meetings have suggested this as an alternative explanation. We consider this possibility with reference to the diagram of cerebellar circuitry in Figure 10.

Antidromic volleys in parallel fibers could excite Golgi cells, which are inhibitory interneurons. However, Golgi cells inhibit the granule cells, which are the source of the parallel fibers. They do not inhibit the Purkinje cells, so this antidromic pathway would appear to be a dead end so far as explaining the conditional pause. We know of no other way that an antidromic signal in a population of directly stimulated parallel fibers could reach a population of unstimulated parallel fibers (see Figure 10). Thus, the hypothesis that there are unstimulated granule cells excited by antidromic signals in the stimulated parallel fibers requires the postulation of an unknown pathway and the postulation of the requisite grandmother-cell dynamics for the unstimulated granule-cells. There is neither evidence nor independent motivation for either postulate.

Parallel fibers also excite stellate cells, which make inhibitory (GABAergic) synapses on the outer reaches of Purkinje cell dendrites. (This effect need not be considered antidromic.) However, potentiation of the parallel-fiber-to-stellate -cell synapse cannot mediate the conditional pause, because the climbing fiber does not innervate the stellate cell. That is, the stellate cell does not have direct access to both the CS and the US signals. This access is essential to the conditioning of an appropriately timed pause. Instead their inputs consist of far fewer parallel fibers and excitation from multiple climbing fibers only through diffusion of spillover glutamate from climbing fibers that contact Purkinje cells (Szapiro & Barbour, 2007). This is directly opposite to the Purkinje cell whose input architecture enables more than 100,000 different parallel fiber inputs to predict one and only one climbing fiber input.

One might, however, entertain the following hypothesis: Parallel fiber stimulation excites a population of inhibitory stellate cells antidromically, orthodromically or both. The excited population consists of temporal grandmother cells with an appropriate range of delayed-peak responses. The climbing fiber stimulation selectively and enduringly enhances the postsynaptic effects of the GABA released onto the Purkinje cell from those stellate grandmother cells whose firing peaks at the time of climbing fiber stimulation. This inhibitory action of GABA is not blocked by doses of gabazine sufficient to block the profound inhibitory effect of off-beam stimulation.

The following considerations weigh against this alternative hypothesis:

- It does not explain why blocking the mGluR7 receptor blocks the conditional pause. The mGluR7 receptor is postsynaptic to the glutamatergic parallel fiber input to the Purkinje cell. It is not postsynaptic to the stellate cell GABAergic input, nor is it a GABA receptor. Thus, on this hypothesis, there appears to be no explanation for the fact that blocking the mGluR7 receptor blocks the conditional pause.
- There is no evidence that the stellate cells have the requisite dynamics.
- A fortiori, there is no evidence that these dynamics are triggered by the first volley in parallel fiber and are unaffected by the temporal distribution of subsequent volleys. Known synaptic mechanisms rarely have the latter property.
- There is no evidence that gabazine, a general-purpose GABA blocker, fails to block the inhibitory effect of the GABA that the stellate cells release onto the Purkinje cells.

In the light of current neuroanatomical, neuropharmacological and electrophysiological evidence, there does not appear to be a plausible alternative to the hypothesis that the mechanisms that time, record and read out the CS-US interval in eyeblink conditioning are intrinsic to the cerebellar Purkinje cell.

### Implications of the Current Results for the Putative Intracellular Engram-Reading Processes

- The pause is created by an abrupt shut down of the mechanism that generates the rapid but extremely irregular spontaneous firing of the Purkinje cell; the pause is not a graded modulation of that endogenous firing rate; it is a complete shutdown, with no measurable sloping off in the firing rate prior to the cessation of firing.
- The shutdown is triggered by the arrival of the first synchronized volley of nerve impulses in a subset of directly stimulated presynaptic parallel fibers.
- Later volleys have no effect on the pause, regardless of their temporal distribution. If there is no cell-intrinsic temporal memory, there has to be a temporal code in the input signal. It seems impossible to reconcile that assumption with the fact that the elicited pause is the same whether stimulation of the parallel fibers lasts 20 milliseconds or several hundred milliseconds and whether the pulse frequency is 50 Hz or 500 Hz.
- When the CS-US interval is short, the latency to shut down of the firing can be shorter than 20 ms; the engram read-out mechanism can shut down firing in less than 20 ms after the synaptic input that triggers read-out.
- The pause onset latency is, however, determined by the mechanism that has encoded the duration of the CS-US interval, because the latency to shut down firing is proportional to the duration of the remembered interval.
- The latency to terminate the shutdown is likewise determined by the engram; it, too, is proportional to the duration of the remembered interval.
- The correlation structure for the pause parameters implies that the two latencies are produced by independent readings of the engram: The offset (↓) and onset (↑) of the shutdown are only weakly and inconsistently correlated (see the ↓v↑ correlations in Figure 9). This stochastic independence has the consequence that the duration of the shutdown (M) is strongly negatively correlated with onset latency (Mv↑ in Figure 9) and strong positively correlated with offset latency (Mv↓ in Figure 9). In other words, a late onset of the pause predicts a short shutdown and an early offset retrodicts a short shutdown.
- The process leading to the initiation of the shutdown and the process leading to its termination are independently initiated by the synaptic input that causes the reading of the engram. If the process leading to termination were initiated by the onset of the shutdown, then the latency to terminate the shutdown would be positively correlated with the latency to initiate it and the CoV of the shutdown latency would be greater than the CoV of the initiation latency. In fact, however, the two latencies are uncorrelated or even weakly and inconsistently negatively correlated (Figure 9), and the CoV of the termination latency is smaller than the CoV of the onset latency (Figure 6).
- The endogenous spike-generating process is not Poisson. The Fano Factors are generally much greater than expected from a stationary Poisson process. The distribution of the endogenously generated interspike intervals has an extremely abrupt rise after a “refractory” interval that ranges from 3 to 8 ms, depending on the cell, an initially steep decline and a remarkably prolonged tail. This tail includes the very long interspike intervals that constitute the conditional pauses. However, when these very long intervals occur at any time other than immediately after CS onset, their duration is not proportional to the CS-US interval, the interval encoded in the engram.
- The greatly prolonged tail in the inter-spike interval distribution of the spontaneously firing Purkinje cell means that pauses like those that constitute the conditional pause appear often. Whatever the process is that produces these spontaneous pauses, the process that produces the CS-conditional pause can preempt or supervene on a spontaneous pause. The unusually long interspike intervals that constitute the conditional pause (most often a single such interval) may begin during one of the spontaneously occurring long interspike intervals. When the CS-US interval is short, this often produces a conditional pause that appears to begin before CS onset.

## A Molecular Biological Agenda

The engram is the mechanism that preserves facts gleaned from experience for use in computations to be performed in the indefinite future. The simplest and most readily varied facts are the quantitative facts. The duration of the interval between two events, such as the onset of a conditional stimulus and the event that it predicts, is an example of an easily varied quantitative fact. A fact of this kind, the duration of the CS-US interval, plays two fundamental roles in associative learning (Balsam et al., 2010; Balsam, Fairhurst, & Gallistel, 2006; Balsam, Drew, & Yang, 2002; Balsam & Gallistel, 2009; Drew, Zupan, Cooke, Couvillon, & Balsam, 2005; Gallistel & Balsam, 2014; Gallistel & Gibbon, 2000; Gibbon & Balsam, 1981; Morè & Jensen, 2014; Ward et al., 2012):

- The duration of the CS-US interval and whether that duration is fixed or variable determines the latency at which the conditional response follows the onset of the CS, and, the pattern of responding within the CS.
- The ratio between the average CS-US interval and the average US-US interval determines trials to acquisition and the vigor of responding during the CS.

Because the Purkinje-cell engram preserves the duration of the CS-US interval that the cell has experienced, there must be an invertible (one-one) mapping between the duration of an experienced interval and the structural change it produces in the engram. In other words, there must be a code (Gallistel, 2017). Because the engram appears to be intrinsic to the Purkinje cell, one may infer that the structural change that encodes the duration of the CS-US interval is a change in a cell-intrinsic, molecular-level structure. Thus, a key step in the discovery of the material realization of the engram must be the deciphering of the code that relates a molecular change within the Purkinje cell to the duration of the inter-event interval that induces it and that it encodes. The discovery of such a mapping would be powerful evidence that the material realization of the engram (or at least an engram) had at long last been discovered. Decisive evidence would come from artificially inducing the structural modification that encoded a given interval and demonstrating that the duration of the firing pause in response to parallel-fiber input is predicted by that modification. This confirmation of the cell-intrinsic molecular-engram hypothesis would be analogous to the eventual confirmation of two hypotheses that were once profoundly controversial: 1) brains contain clocks (Beiling, 1929; Richter, 1922; Wahl, 1933); 2) the clock is a molecular-level cell-intrinsic mechanism that does not depend on neural circuitry for its function (Ralph, Foster, Davis & Menaker, 1990; Le Sauter & Silver, 1994; Silver, Le Sauter, Tresco & Lehman 1996; Bruce & Pittendrigh, 1956). Neuroscientists have been slow to recognize the role of molecular level cell-intrinsic mechanisms in the complex computations that are the foundation of cognition.

The search for the molecular engram in the Purkinje cell should probably begin with a focus on the metabotropic mGlu7 receptor, because blocking that receptor prevents presynaptic input from the parallel fibers from triggering the conditional pause. Metabotropic receptors convert extracellular signals (the transmitter release by a presynaptic spike) into intracellular biochemical cascades. The first step in most such cascades is a structural change in a G protein. G-protein-initiated cascades are known to lead to alterations in many aspects of cellular physiology, including gene transcription. In this case, the cascade must read the engram before it comes to the membrane-intrinsic ionic channel or channels whose modification abruptly shuts down endogenous firing, because the latency to the shutting down of the endogenous firing is determined by the encoded duration. Given this knowledge and the further facts revealed by this analysis, it seems that a concerted investigation of this intracellular biochemical cascade could be expected to bear fruit within a reasonable time span.

## Acknowledgements.

This work was supported by grants from the Swedish Research Council to G.H. (2018-03191) and to F.J. (2019-02034), from the Åke Wiberg Foundation and the Magnus Bergvall Foundation to F.J., from Åke-Wiberg Foundation (M19-0375) and the Segerfalk Foundation (2019-2246) to A.R., and by NSF Graduate Research Fellowship, Award no. 1644760 to M.R.

